# SmartScope2: Simultaneous Imaging and Reconstruction of Neuronal Morphology

**DOI:** 10.1101/118927

**Authors:** Brian Long, Zhi Zhou, Ali Cetin, Jonathan Ting, Ryder Gwinn, Bosiljka Tasic, Tanya Daigle, Ed Lein, Hongkui Zeng, Peter Saggau, Michael Hawrylycz, Hanchuan Peng

## Abstract

Quantitative analysis of neuronal morphology is critical in cell type classification and for deciphering how structure gives rise to function in the brain. Most current approaches to imaging and tracing neuronal 3D morphology are data intensive. We introduce SmartScope2, the first open source, automated neuron reconstruction machine integrating online image analysis with automated multiphoton imaging. SmartScope2 takes advantage of a neuron’s sparse morphology to improve imaging speed and reduce image data stored, transferred and analyzed. We show that SmartScope2 is able to produce the complex 3D morphology of human and mouse cortical neurons with six-fold reduction in image data requirements and three times the imaging speed compared to conventional methods.

## Introduction

The forefront of bioimaging technology is often driven by a desire for faster collection of larger image volumes, with increased spatial and temporal resolution. This has led to numerous technological developments, including great strides in optical engineering, new optical techniques such as multiphoton fluorescence microscopy^1^, light-sheet imaging^2^, and several modes of super-resolution microscopy^3^. This trend creates large and complex biological image data sets, leading to ongoing challenges in image visualization and analysis^4^. As these computational challenges are encountered and overcome, it’s worth recalling that imaging is not an end in itself, but is rather a means for obtaining data and scientific understanding from biological systems. In this context, the integration of computational processing with automated microscopy methods has the potential to allow efficient collection of optimal datasets in complex imaging experiments with far less hands-on involvement than in the past. One recent example of research in this direction is adaptive optimization of light sheet microscopy^5^, which improves light sheet imaging by automatically determining the imaging parameters over time to yield consistent, optimal image quality despite spatially and temporally variable imaging conditions. Additionally, recent improvements in acousto-optical deflector (AOD) instrumentation^6–8^ have enabled continuing development of rapid, random-access two-photon microscopes. The speed and flexibility of AOD systems has provided a useful avenue for online motion-correction and imaging fluorescent activity sensors from arbitrary scan volumes. Other researchers have recognized the importance of these capabilities, and similar efforts to optimize image data collection are being made without AOD technology^9,10^. Despite the ongoing interest in adaptive imaging experiments, we are unaware of any existing approaches to apply online computational processing to an important factor in neuronal cell type identification: imaging and reconstruction of neuronal morphology^11^.

The morphology of neurons is often sparse; the neuron occupies only a small fraction of the total volume explored by its dendrites and axons. Because of this sparsity, imaging the total volume bounding a neuron is highly inefficient. Additionally, the topology of all neurites emanating from the cell body provides a means to access and trace the full morphology starting from the soma. Consequently, we set about to use the neuronal structure itself to guide image collection and extract neuronal morphology without imaging the large volumes devoid of signal. In this paradigm, both image data and imaging time are reduced by tailoring the spatial distribution of imaging areas to the sample through on-line processing.

Our system, SmartScope2 (denoted as “S2” below for simplicity), is illustrated in Fig. 1. S2 is an open-source software module that combines control of a two-photon (2p) microscope with simultaneous analysis of neuronal morphology in 3D image volumes, following the similar concept of automated analysis-based imaging on the confocal SmartScope system^12^. The software is a plugin for Vaa3D^13^, an open-source, cross-platform application for image visualization and analysis of 3D image data. We designed S2 to communicate with a commercial 2p microscope via transmission control protocol / internet protocol (TCP/IP) connections, enabling S2 to be operated remotely and flexibly to analyze image data transferred from the microscope to an S2 client workstation. Remote operation of S2 also removes the computational load of neural reconstruction from the data acquisition microscope, and thus could facilitate cloud-based automated microscopy and morphological reconstruction tasks.

**Figure 1.**
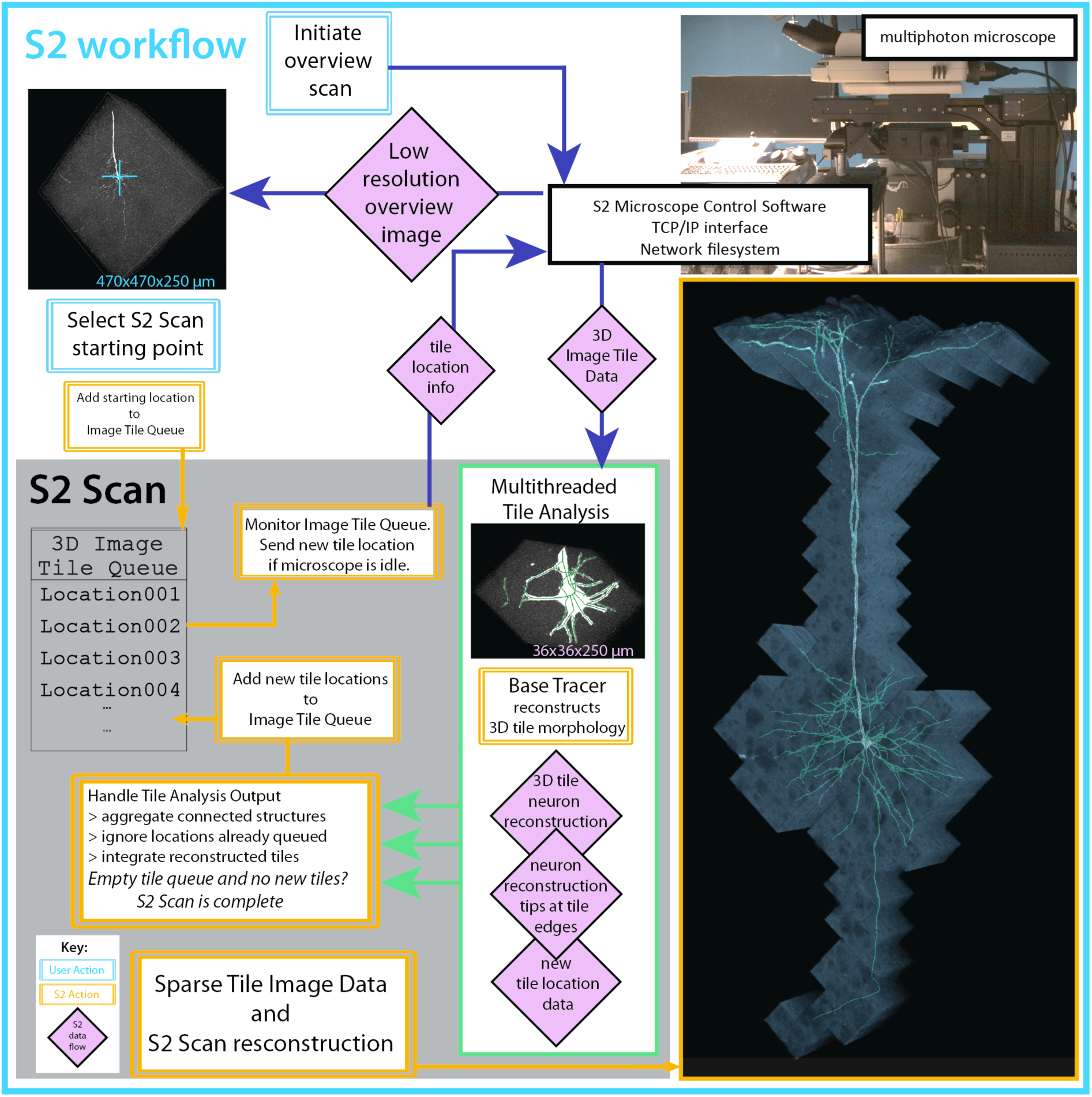
S2 schematic and workflow. S2 operation starts by collecting a low-resolution overview stack covering the full scan field of the microscope. After the user selection of cell-bodies using single-click ‘virtual finger’ selection in Vaa3D, the cell body becomes the starting point for the S2 scan process and the first tile added to the 3D Image Tile Queue. S2 constantly monitors the queue, and if the microscope is not imaging and there is a location in the queue, a 3D image stack is initiated at that location. Image data from the microscope is traced using a user-chosen tracing method, automatically generating a 3D reconstruction of the neuron within the image tile. If this reconstruction contains any elements near the tile borders, those tip locations are used to define a new tile location. This tile location (with accompanying tip locations) is passed to the 3D Image Tile Queue. This process continues until all tiles have been imaged and processed. The lower right panel is a composite illustration of the alpha-projection, maximum-intensity projection and the S2 reconstruction of S2 Scan 2 in Table S1.

To reconstruct 3D neuronal morphology with minimal imaging, we utilized the S2 scan strategy (Fig. 1, Methods). First, a low-resolution overview scan is collected where a target neuron cell-body is chosen by the user (or determined automatically using existing algorithms^14^). Second, a 3D image stack (“tile”) centered on the target cell body is scanned and an automatic tracing algorithm (the “base tracer”) is applied to that tile data. Third, the resulting 3D digital reconstruction is analyzed to determine if any tips of the neuron structure extend to the edges or corners of the tile. Fourth, the next tile containing those tip locations is imaged and the 3D reconstruction process is continued in the next tile. The imaging and reconstruction processes are iterated until all traced structures have terminated within an imaged tile. The process is automatic, proceeding until all of the neuron’s morphology in the sample captured by the base tracer has been reconstructed.

Instead of implementing the S2 scan in a serial fashion where image collection and reconstruction alternate in time, our software design decouples image collection from image analysis to minimize the impact of computational time on the total S2 scan time. Specifically, any new tile locations generated from analysis of a 3D tile are added to a queue of tile locations to be scanned, and S2 monitors this queue and initiates imaging a new tile whenever the microscope is not imaging. This design, combined with dedicated threads for image analysis, minimizes the idle time of the microscope.

We first tested S2 on individual neurons, simultaneously producing sparse volumetric image data and neuronal reconstructions. These scans included EGFP-expressing neurons from mouse cortex (Fig. 2A, Fig. 2B, Fig. S1A, Table S1 scans 1-3). The reconstructions show tracing of dendrites, including the basal skirt and apical tuft of a deep-layer pyramidal cell, as well as descending axons of genetically-labeled mouse neurons. These S2 scans resulted in an average of 4.6-fold reduction in imaged volume compared to a rectangular bounding box surrounding the same structure (N = 3 individual neurons, S2 scans 1-3 in Table S1). The summed area of all tiles, including a user-defined 10% overlap between adjacent tiles, was also reduced by 4.0-fold relative to the bounding box approach. This reduction in image area can be regarded as a conservative estimation of S2’s ability to reduce imaging load for neuronal reconstruction. To define this rectangular bounding box without S2, either a human operator would have to manually follow dendrites from the cell body until they terminate or leave the sample, or some automated method would have to be deployed to identify the ends of all of the connected structures. The latter choice was exactly the stopping criteria for the S2 scan, and we are unaware of any other methods that accomplish this. Therefore, a more realistic comparison is to use an “integer bounding box” made of an integer number of tiles that are the maximum size for our microscope at this resolution. This comparison reveals scan area reductions of eightfold for the EGFP-expressing cortical neurons (8.26, N = 3) (Table S1, scans 1-3). This improvement in scan area also results in a reduction of image data averaging 7.1-fold, which includes the overlap between adjacent tiles (Table S1, “Area Efficiency”).

**Figure 2.**
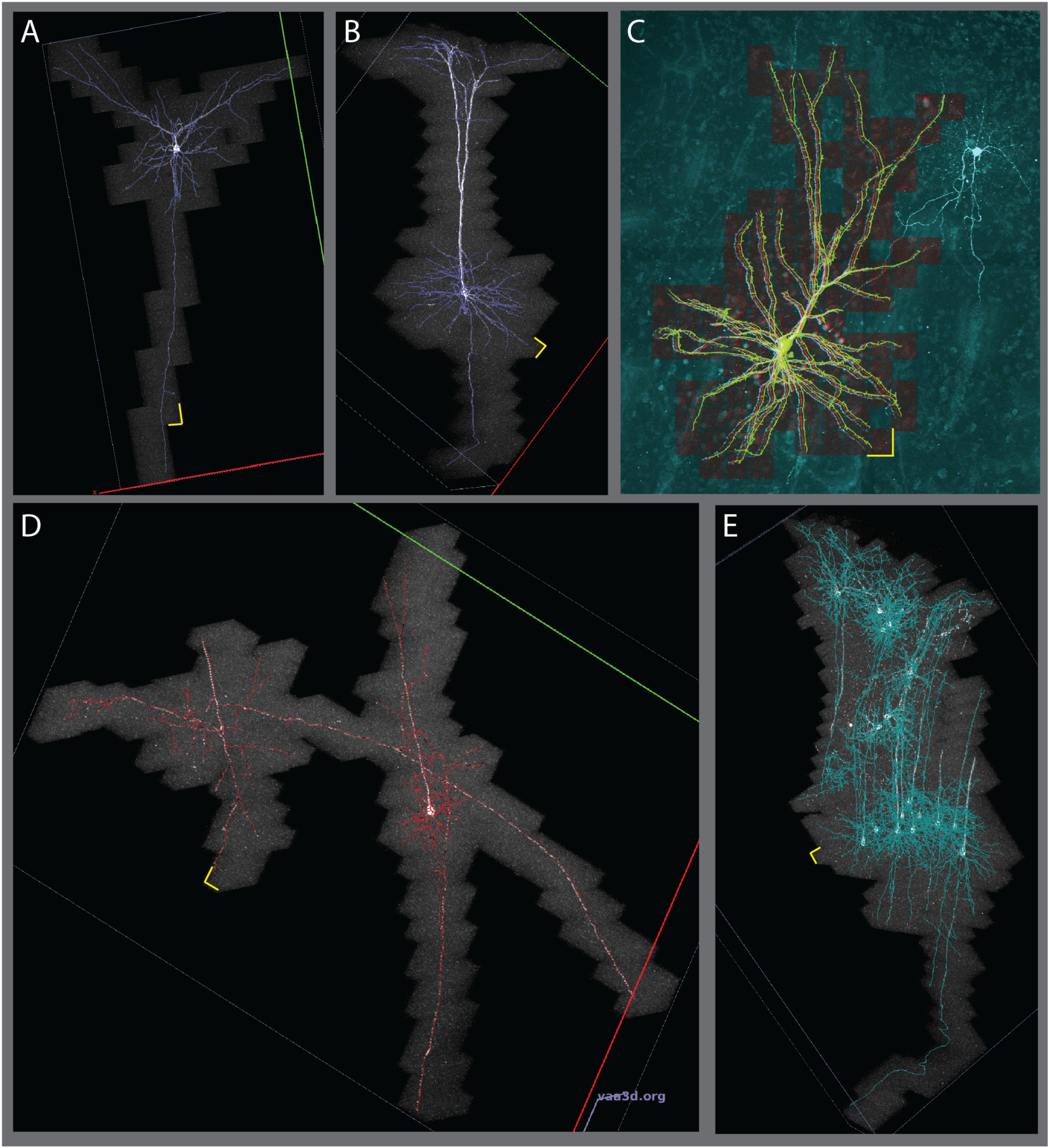
S2 scans showing sparse imaging of cortical neurons. A. A layer 2/3 pyramidal cell, with extensive tracing of descending axon. B. Deep-layer pyramidal neuron in mouse successfully reconstructing apical tufts, basal dendrites and descending axon. C. S2 scan of a human cortical pyramidal neuron filled with Alexa 488 during whole-cell recording. S2 scan tiles (red) and S2 reconstruction (green) are overlaid on a traditional raster-scanned image (cyan). The S2 scan successfully traced continuous signal in apical and basal dendrites without crossing over to trace processes from the nearby labeled aspiny cell. D. S2 scan in mouse cortex initiated at the soma of the deep-layer cell on the right. S2 followed the dendrites of this cell onto nearby axons from two other cells. E. S2 scan in mouse cortex with relatively dense labeling. S2 followed dendrites and axons of several cells spanning the depth of cortex. Yellow scale bars are 36 μm.

The reduction in scan area and image data we achieved suggests S2 scan could also shorten imaging time. In fact, the S2 scans on the EGFP-expressing mouse neurons were accomplished an average of 3.5 times faster than the time to image the integer bounding box with the same input laser power. Ideally, the imaging time should improve by exactly the same factor as the image data, but in practice this is not the case due to three factors. First, the effective imaging time per pixel is not exactly constant across image tile sizes. For example, if there were a fixed time overhead per line (as might be the case for unidirectional non-resonant galvo scanning) or per plane (e.g. due to fixed settling time of z movement), the effect of these fixed times would be proportionally greater for smaller tiles. We observed this effect in a test consisting of N = 20 sparsely-labeled neurons in 50-μm sections (Fig. S3 and Supplemental Text). For the current data set in Fig. 2 and Table S1, we used bidirectional resonant scanning and zero-delay piezo z-actuator movement to eliminate these effects. Second, any software or hardware overhead per tile can easily accrue into a substantial time cost as the number of tiles becomes large. Third, the computational time associated with reconstructing each tile can affect the total imaging time, especially if the computation is serially interleaved with imaging. Our strategy of queuing locations to avoid microscope idle time is key to minimizing this factor. For EGFP-expressing mouse neurons, single-thread computational analysis time was 77% of imaging time, thus a simple serially interleaved imaging and analysis paradigm would require 1.77 fold longer than the imaging time alone. However, our optimization of asynchronous imaging and parallel processing reduced the effect of analysis time such that S2 scans took only 9.5% longer than the total imaging time.

In addition to imaging and reconstructing mouse cortical neurons, we applied the S2 system to a human neuron filled with a fluorescent dye in an acute slice preparation from re-sectioned, surgically-excised neocortical tissue (Fig. 2C, Table S1 Scan 4). Imaging and reconstruction of fluorescently labeled cells in adult human cortical tissue can be challenging because of the presence of auto-fluorescent lipofuscin granules in neuronal cell bodies. The broad emission spectrum of lipofuscin spans green and red emission bands, manifesting as bright puncta in both red and green channels. Before S2 scanning in human tissue, we tuned the red channel PMT gain so mean intensity values from lipofuscin fluorescence were equal in both red and green channels. We then implemented an on-line intensity subtraction so the S2 base tracer operated on a linear unmixed channel G-αR, reducing the impact of lipofuscin fluorescence on S2 auto-tracing of Alexa 488-filled cells. For the human neuron data presented here, *α* = 1.0, but the parameter is user-adjustable in the S2 user interface. For the layer 2 pyramidal neuron in S2 Scan 4 (Fig. 2C), the dendritic morphology was relatively compact, but the sum of the S2 tiles was still 4.0-fold less area than the integer bounding box. The combined human and mouse neurons (S2 scans 1-4 in Supplemental Table S1) have an average area efficiency of 6.35 and average speed improvement of 3.1 compared to the integer bounding box.

We used two approaches to evaluate the reconstruction accuracy of S2. First, for three isolated EGFP-expressing neurons in mouse (Supplemental Table S1, Scans 1-3), we compared S2 to manual reconstructions generated by three annotators who traced the neuron structure manually using Vaa3D-Terafly^15^. We compared the similarity between the S2 reconstruction and the manual reconstructions as well as the similarity among manual reconstructions, using the best average spatial distance score *d*^16^ (Supplemental Text and Table S2). Overall, the average bi-directional distance scores between S2 and three manual reconstructions are very close to the respective scores between three pairs of manual reconstructions: *d*_S2-manual_ = {5.74, 8.62, 4.09} and *d*_manual-manual_ = {5.25, 5.42, 3.63} voxels in scans {1, 2, 3}. In scan 2, the high bi-directional score was actually due to a high average directional distance score from S2 to manual (13.42 voxels vs 3.83 voxels) on average, indicating that in that particular case, S2 reconstructions actually captured more neurite segments than human annotators. We applied our second approach to verify S2 accuracy to the human neuron shown in Fig. 2C. Here, we collected four large tiles of image data encompassing the entire labeled structure and overlaid the S2 scan and its reconstruction onto the tiled imaged, as seen in Fig. 2C. The resulting image shows that S2 reconstruction may not trace the morphology at the distal tips of dendrites that exhibit some beading (Fig. S6), but accurately includes all the continuously labeled dendrites, while avoiding the neurites of a nearby aspiny neuron that was also filled in this sample.

After demonstrating S2 scanning of individual neurons, we also targeted S2 to areas with structures from multiple labeled neurons in close proximity (Fig. 2D and E, Fig. S1 B). Such samples could be generated by sparse Cre recombination in a fluorescent protein reporter mouse line (as here), from multi-patch experiments in acute brain slices or from sparse viral labeling in cultured brain slices. The current version of S2 will image and connect all structures that are contiguous with structures traced in the first tile. In this way, labeled neurons with closely apposed neurites will all be imaged and traced by S2. In Fig. 2D, initiating tracing on a pyramidal cell led to additional tracing of an ascending axon and a collateral of a descending axon from a neighboring neuron whose cell body was in an adjacent slice. The extremely sparse nature of these neurites led to an increased benefit of S2 scanning compared to conventional imaging: 8.5 times smaller data size and 3.7 times faster acquisition than a comparable bounding box scan. Targeting other multi-neuron samples led to large datasets spanning up to 2 mm and including up to 19 neurons. Even for multi-neuron scans, the output of S2 is a connected structure by design, which could potentially be dissected into separate neuron trees post-hoc using analysis methods targeted specifically to this problem^17^. Using S2 to image multi-neuron samples is appealing for its automatic reconstruction of multiple neurons, but may be limited in improving acquisition speed because some samples with multiple neurons (e.g. S2 scans 5 and 6 in Table S1.) have less dramatic sparsity than individual neurons.

## Discussion

S2 provides a means of rapidly imaging and tracing morphology of fluorescent neurons in sparsely-labeled samples. S2 is currently implemented with a multiphoton microscope designed for imaging of fixed samples up to ~350 μm thick, but the S2 concept could in principle be extended to any imaging modality where data collection can be spatially localized within a sample. AOD-based random-access microscopes are natural targets for implementing future versions of S2 because the truly random-access imaging of AODs can allow for fast data acquisition in arbitrary volumes. Also, a stage-based scanning approach such as the one used to collect the data in Fig. 2 can compensate for the relatively limited scanning fields of view associated with current AOD systems. The sample slice thickness used here (which is typical for electrophysiology) is expected to result in some loss of dendritic and axonal arbors due to truncation by physical sectioning. Previous work quantifying these losses suggests that rat layer 2/3 pyramidal cells can expect 89% complete dendrites and 60% complete local axonal arbors in 300 μm-thick samples^18^. While thicker samples could enable marginal gains in the completeness of local structures, the ideal neuronal reconstruction would include the entire dendritic and axonal arbors that can extend to numerous brain areas and across several millimeters in the mouse. Imaging across millimeter length scales while maintaining the high resolution needed to reconstruct neuronal morphology typically implies terabytes of image data, most of which contains no signal from the neurons of interest. Existing terabyte-scale datasets can be processed using specialized analysis tools such as the recently developed Ultratracer^19^, which shares the same underlying operational philosophy as S2. One avenue to efficiently image entire neurons in larger samples would be to combine S2 with random access large-scale imaging techniques such as tiled SPIM^20^ to achieve whole-neuron reconstructions while imaging only a fraction of the sample volume.

S2 provides a platform for further development of global analysis of neuronal structure and imaging parameters in an online manner. Furthermore, this approach may also improve efficiency in other imaging modalities where sparse, continuous structures can automatically guide data collection. The modular implementation of S2 that remotely connects software user interface, data storage, data analysis, and hardware design together will allow more sophisticated smart imaging paradigms to be established based on S2.

## METHODS

### Microscope Hardware and Configuration

The SmartScope2 system was implemented using an upright resonant-scanning two-photon microscope (Investigator, Bruker FM) with one resonant galvanometer mirror and two non-resonant galvanometer mirrors in series. Imaging was performed using a tunable pulsed laser (Coherent Chameleon Ultra II) at 920 nm and intensity-controlled with a Pockels cell. To maximize fluorescence collection from tissue slices, the microscope was configured with both epi-and trans-detectors, including a high-numerical aperture oil condenser to maximize trans-collection efficiency. The S2 hardware includes a 400 μm-travel piezo z-actuator for the objective lens (Nikon 16x water immersion, NA = 0.8, or Olympus 25x water immersion, NA = 1.05 which was used for the scans shown here), allowing rapid, resonant-scanned z-stacks to be positioned at any location within the field of view. It is able to rapidly image fluorescence signal over the entire depth of acute brain slices (typically 300~350 μm thick), scan both small volumes (~36×36×300 μm) and large areas in resonant mode (474 × 474 μm for the 25x, 800x800 for 16x), and allow simple, remote control of high-level microscope functions. The ability to position a resonant-scanned 3D region-of-interest (ROI) at arbitrary locations using either galvanometer steering or stage movements allows us to execute the S2 scan within the microscope field-of-view (FoV) or to follow neuron morphology laterally beyond a single field of view to the millimeter scale, practically limited only by the size of the sample.

### S2 Software and Code Availability

The S2 microscope control module was written in C++ as a plugin to Vaa3D. S2 source is available at https://github.com/Vaa3D/vaa3dtools/tree/master/hackathon/brl/s2 In addition to being available as source, the S2 plugin will be included as a plugin in future binary releases of Vaa3D. Currently, the user can select between three tracing methods, APP2^14^, MOST^21^, and Neutube^22^ The data collected here utilized Neutube as the base tracer.

The low-level microscope control (e.g. scan voltage waveforms, digital-to-analog conversion, etc) was handled by alpha-release microscope control software from Bruker (PrairieView version 5.4 alpha), which allows high-level control of the microscope via TCP/IP connection. Using two dedicated TCP/IP connections, the microscope was polled for current state information over one connection while a separate connection was reserved for sending commands to initiate image tile collection at specified locations. In principle, the S2 software design allows for this approach to be extended to other microscopes, provided that the microscope can be externally controlled through C++ code.

### Imaging Parameters

Overview scans are collected at 1x optical zoom (512×512 pixels) over the entire sample depth with 20 micron z-steps. Tile scans were typically collected at 13x zoom (the highest stable resonant zoom available on our system) with a voxel size of 0.23 × 0.23 x 1.0 μm to sample the 25x objective point spread function at roughly twice the full-width-at-half-maximum (FWHM) values along x, y and z. S2 tiles were imaged without averaging, but under these imaging parameters the acquisition software uses 11 samples from the photomultiplier tube readout to generate each of the 157 pixels per line. The time for maximum-sized tiles (1.18x optical zoom, 1730×1730 pixels) was determined by imaging at the same laser power as the S2 tiles and averaging 8 times frame-by-frame to emulate the 11x multisampling of the S2 image tiles (8x averaging is the largest value available in software without exceeding the 11x multisampling). This approach maintains the approximate excitation dose per pixel with constant laser power. Individual target neurons were selected visually for brightness, local sparsity of labeling and completeness of the morphology within the slice.

### Mouse neuron labeling

Ai139 mice (manuscript in preparation) were injected with an AAV viral vector that generates sparse Cre recombination (manuscript in preparation) using a Nanoject II microinjector (Drummond Scientific, Broomall, PA) within neocortex. Four weeks following injections animals were perfused and 300 um slices were prepared for imaging.

### Human *ex vivo* brain slices

The human neocortical tissue specimen was obtained during brain surgery for a patient with intractable epilepsy. It was necessary to remove the overlying neocortical tissue to gain access to the underlying diseased tissue. Informed patient consent was obtained for use of neocortical tissue for research purposes under a protocol approved by the institutional review board of Swedish Medical Center (Seattle, WA). Brain slices of 350 μm thickness, spanning the majority of a gyrus and in a plane perpendicular to the pial surface were prepared using a Compresstome VF200 (Precisionary Instruments) using the previously described NMDG protective recovery method^23^.

### Single human neuron cell filling via patch clamp recording

Human neocortical pyramidal neurons were targeted for whole-cell patch clamp recording using infrared differential interference contrast optics and a 0.8 NA 40X water-immersion objective (Olympus America) and were filled intracellularly by inclusion of a saturating concentration of Alexa 488 hydrazide in the patch pipette. The intracellular pipette solution composition was (in mM): 130 K-Gluconate, 4 KCl, 10 HEPES, 0.3 EGTA, 10 phosphocreatine-Na_2_, 4 MgATP, 0.3 Na_2_-GTP. The pH was adjusted to 7.35 with 1M KOH and the osmolality is adjusted to 285-290 mOsm/Kg using sucrose as needed. The duration of whole cell recording was 15-20 minutes in order strike the best compromise between achieving completeness of cell filling and preserving neuron health and membrane integrity. Upon termination of the recording session, the human brain slices were fixed in 4% PFA for ^~^20 hours and then transferred into 1X PBS plus 0.05% sodium azide for storage prior to imaging. Human cortical tissue was obtained from patients undergoing surgery to treat medically refractory epilepsy. Patients underwent standard surgery for temporal lobe epilepsy and provided informed consent to have their tissue studied under an IRB approved protocol. All samples were mounted in Fluormount G (Southern Biotech) before imaging.

### Analysis and Statistics

To characterize the ability of S2 to track structures across tiles, we developed the S2 quality index *Q_S2_* given by *Q_S2_* = 1 − (*N_edge tips_ – N_tips_f_edge_*)*/N_tips_* Where *f_edge_* is the fraction of tile area considered edge - here we use the area within 5% of each edge, totaling 19% of the tile area, so 19% of randomly distributed tips are expected to be identified as edge tips purely by chance. *N_tips_* is the number of terminal tips of a single reconstructed neuron tree. Within *N_tips_*, *N*_edge tips_ is the number of terminal tips located in the edge area. The edge tips in excess of the expected number are presumed to result from failures to continue tracing across tile boundaries, so a scan with no excess edge tips will have a quality index of 1.0 while a scan with three times as many edge tips as expected would result in quality index of 0.62.

All average values for S2 performance are from three individual mouse neurons (S2 scans 1 and 2 shown in Fig. 2 A and B, and scan 3 in Figure S1), with the complete information for each scan displayed in Table S1.

## Acknowledgements

We thank Hsien-Chi Kuo for manual neuron reconstruction, Shenqin Yao and Ben Ouellette for preparing the sparse neuronal expression sections, and Christof Koch for discussions. The authors wish to thank the Allen Institute founders, Paul G. Allen and Jody Allen, for their vision, encouragement and support.

